# *ct2vl*: Converting Ct Values to Viral Loads for SARS-CoV-2 RT-qPCR Test Results

**DOI:** 10.1101/2022.06.20.496929

**Authors:** Elliot D. Hill, Fazilet Yilmaz, Cody Callahan, Annie Cheng, Jasper Braun, Ramy Arnaout

## Abstract

RT-qPCR is the de facto reference method for detecting the presence of SARS-CoV-2 genomic material in infected individuals (1). Although RT-qPCR is inherently quantitative and despite SARS-CoV-2 viral loads varying by 10 orders of magnitude and therefore being potentially highly clinically informative, in practice SARS-CoV-2 RT-qPCR results are usually reported qualitatively as simply positive or negative. This is both because of the mathematical complexity of converting from *C*_*t*_ values to viral loads and because the same *C*_*t*_ value can correspond to orders-of-magnitude differences in viral load depending on the testing platform (2, 3, 4). To address this problem, here we present *ct2vl*, a Python package designed to help individual clinical laboratories, investigators, and test developers convert from *C*_*t*_ values to viral loads on their own platforms, using only the data generated during validation of those platforms. It allows any user to convert *C*_*t*_ values to viral loads and is readily applicable to other RT-qPCR tests. *ct2vl* is open source, has 100% code coverage, and is freely available via the Python Package Index ([PyPI](https://pypi.org/project/ct2vl/)).

**IMPORTANCE:** Up to now, COVID-19 test results have been reported as positive vs. negative, even though “positive” can mean anywhere from 1 copy of SARS-CoV-2 virus per milliliter of transport media to over 1 *billion* copies/mL, with attendant clinical consequences. Democratizing access to this quantitative data is the first step toward its eventual incorporation into test development, the research literature, and clinical care.

## INTRODUCTION

The real-time reverse-transcription polymerase chain reaction, commonly known as quantitative RT-PCR or RT-qPCR, is a standard method for testing human samples for the presence of viruses such as HIV-1 (human immunodeficiency virus type 1), HCV (hepatitis C virus), and, since 2020, SARS-CoV-2 (5, 6). In RT-qPCR, the tiny amount of genetic material originally present in a positive patient sample is copied by a polymerase enzyme over repeat cycles, resulting in exponential amplification that eventually leads to detectable amounts of genetic material (7). The cycle number at which the detection threshold is reached is called the *C*_*t*_ value. Because the reaction is monitored continuously, the threshold may be crossed between cycles, leading to the alternative term fractional cycle number [FCN] (8). The more starting material, the fewer cycles are needed for signal to cross the threshold. Thus, the smaller the *C*_*t*_ value, the greater the amount of starting material.

RT-qPCR results can be reported either qualitatively (positive/negative) or quantitatively. For most infections, quantitative results are usually reported not as a *C*_*t*_ value but as a viral load: the number of copies of viral genomic material present per unit volume of sample (i.e., a concentration). The most common unit is copies/mL. An important advantage of viral loads over *C*_*t*_ values is that viral loads are consistent across platforms. *C*_*t*_ values are not, due to platform-specific differences in polymerase, am-plication conditions, the signal-detection method, whether “dark cycles” are run, and other factors. For example, for SARS-CoV-2, a *C*_*t*_ value of 26 corresponds to a viral load of 100 copies/mL of viral transport media on one FDA-approved platform and nearly 500,000 copies/mL on another (9). This platform-to-platform variability can make *C*_*t*_ values difficult to interpret and can lead to mistaken conclusions about a patient’s clinical course (Fig. 1). An additional advantage is the direct correlation between disease burden and viral load, as opposed to the “golf score” inverse correlation with *C*_*t*_.

**FIG 1.**
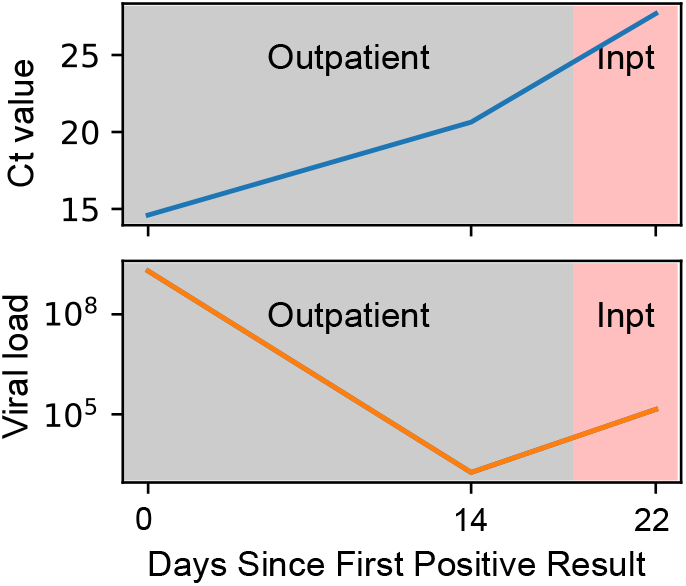
*C*_*t*_ value vs. viral load (in copies/mL). (a) In this patient, *C*_*t*_ values trended consistently upward. Clinically, it would be reasonable to interpret this trend as reflecting a continuous fall in viral load and thereby expect clinical improvement, all other things being equal. Yet the patient worsened between Day 14 and 22, necessitating hospitalization. It happened that the Day 14 result was from a different platform from the Day 0 and Day 22 results; the *C*_*t*_ value for Day 14 cannot be interpreted on the same scale as the other two datapoints; the plot is misleading. (b) Conversion to viral load shows the true picture: a rebound in viral load, consistent with the worsening clinical picture.

In the scramble to create, approve, and validate tests for COVID-19, most SARS-CoV-2 RT-qPCR tests were not validated to output viral load (10). Yet viral loads can be valuable indicators of where a patient is in the course of infection, as well as the likelihood of being infectious. For these reasons, it has been our experience that clinical staff often ask laboratorians informally to report *C*_*t*_ values for their patients, outside of the health record. The difficulty in interpreting *C*_*t*_ values has persisted into the third year of the pandemic. As time goes on, the likelihood that a patient or caregiver will encounter RT-qPCR results from platforms with disparate *C*_*t*_ -value scales cannot but increase, increasing the chance of diagnostic error.

Fortunately, the correspondence between *C*_*t*_ value and viral load in RT-qPCR is well understood mathematically, and the validation studies that laboratories must perform in order to bring a test online can provide the data necessary to convert from *C*_*t*_ to viral load on clinical samples (11). In past work, we wrote computer code to convert from *C*_*t*_ values to viral loads to help reveal and quantify the range of viral burden in the patient population (12) and to compare the sensitivity and utility of testing from different anatomical sources quantitatively and generalizably (e.g. saliva vs. nasal secretions vs. nasopharyngeal secretions) (13, 9). We later expanded on this code to provide viral loads for new platforms brought online at our institution. However, this code applied only to our own platforms, despite SARS-CoV-2 viral loads being a global need. To address this need, here we present a much-expanded new Python package called *ct2vl* intended to make it straightforward to convert from *C*_*t*_ values to viral loads on any platform.

## METHODS

### Mathematical derivation

Traditionally, conversion from *C*_*t*_ values to viral loads has required first creating a standard curve spanning a range of viral loads at least as large as what is observed in clinical practice. However, standard curves can be time consuming and expensive, especially when viral loads range over as many orders of magnitude as they do with SARS-CoV-2 (≥ 1 billion-fold between the lowest and highest viral loads encountered in clinical practice) (12). Fortunately, reliable *C*_*t*_ -to-viral load conversion can also be performed mathematically based on the well understood biochemical principles of PCR (14, 11, 8). This mathematical approach requires only (1) time series of signal vs. cycle number for positive samples and (2) an anchor point— the *C*_*t*_ value for a given viral load—such as labs routinely measure, in replicate, when validating the limit of detection (LoD) before bringing a platform online.

Both the equation we used and its experimental validation have been described in detail in previous work (12), so we review the approach only briefly here. PCR generally exhibits three phases: a lag phase set by the stochasticity of polymerase first encountering template molecules (and in practice the platform’s detection threshold), an exponential (or “log”) phase during which the amount of product roughly doubles each cycle, and finally a plateau phase due to inhibition of the enzyme by the (now-copious) product it has produced. A detection threshold is crossed during the exponential phase; in fact at least one large diagnostics company determines the threshold, and thereby the *C*_*t*_ or FCN, from the cycle at which maximum exponential growth is observed (8). Absent additional considerations, converting from *C*_*t*_ to viral load would involve simply fitting this relationship using an S-shaped function (e.g. Gompertz’ growth curve).

In practice, fitting S-shaped curves is not straightforward. A good fit requires careful weighting of datapoints in different parts of the curve, and in practice no S-shaped function—Gompertz, sigmoid, logistic, or Chapman—precisely captures the part of the curve that is the most important for viral load determination, the exponential phase (15). Moreover, there is no guarantee that the details of the PCR formulation (e.g., multiple targets; internal controls) will not affect the details of the fit. Therefore, an alternative approach is to fit only the key determinant of the region of interest: the decrease in maximum polymerase efficiency that is frequently observed with increasing cycle number, especially in situations where template is admixed with an internal-control template whose amplification can compete with the template for polymerase, and inhibit polymerase, in more complex ways than a model of a single template might capture.

Exponential growth with decreasing replication rate yields the following equation for viral load, *v*_0_, as a function of *C*_*t*_ value (see the supplementary information of Arnaout et al., 2021 for the derivation):

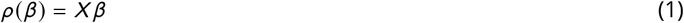

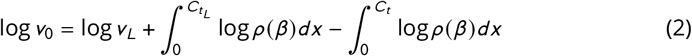

Here, *v*_*L*_ and 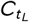 give the anchor point: the simplest anchor point is to let *v*_*L*_ be the limit of detection (LoD) (12) and 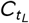 be the *C*_*t*_ value at the LoD. *ρ* is a polynomial fit of maximum replication rate vs. cycle at maximum replication rate (a slight change from (12)). These constitute the parameters of this model (see Implementation, below). Maximum replication rate and the cycle at maximum replication rate are derived from time series data of the form amount of material vs. cycle number (Fig. 2a). The amount of material is most often measured as a fluorescence intensity (e.g. of an intercalating fluorophore present in the RT-qPCR reaction mix).

**FIG 2.**
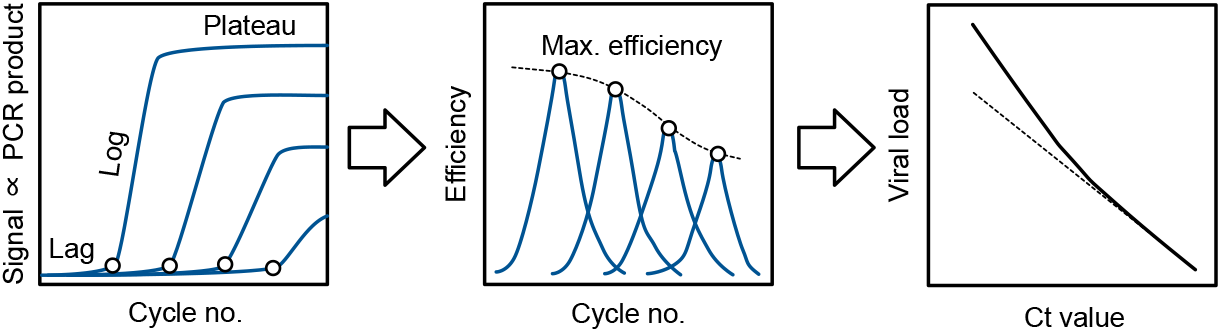
Overview of *ct2vl*’s approach. Left: Signal vs. cycle number traces are analyzed to find maximum replication efficiency (open circles). Middle: The fall in maximum efficiency vs. cycle number is fit by a curve (dotted line). Right: Together with an anchor point (Ct at the LoD), this curve is used to convert Ct values to viral loads (solid line). Without this curve, viral load would be underestimated (dotted line).

### Implementation

To parameterize Equation (1), we must find the coefficients, *β*, of the polynomial regression fit between the maximum replication rate and the cycle at maximum replication rate. To calculate max replication rate and cycle at max replication rate, a set of PCR traces for positive samples are obtained from the platform and processed as follows. The initial 3 cycles are removed because these initial values of PCR traces are often noisy and may interfere with the estimation of maximum replication rate. Negative signal-intensity values are considered noise and therefore set to 0. The data is smoothed, ensuring monotonic increase (PCR product cannot decrease; slight/transient decreases sometimes observed during the lag phase are attributed to signal-detection noise). These steps result in denoised traces. Examples of processed traces are plotted in Fig. 3a. Even after denoising, some noisy measurements of replication rate can be observed in the early cycles (Fig. 3b).

**FIG 3.**
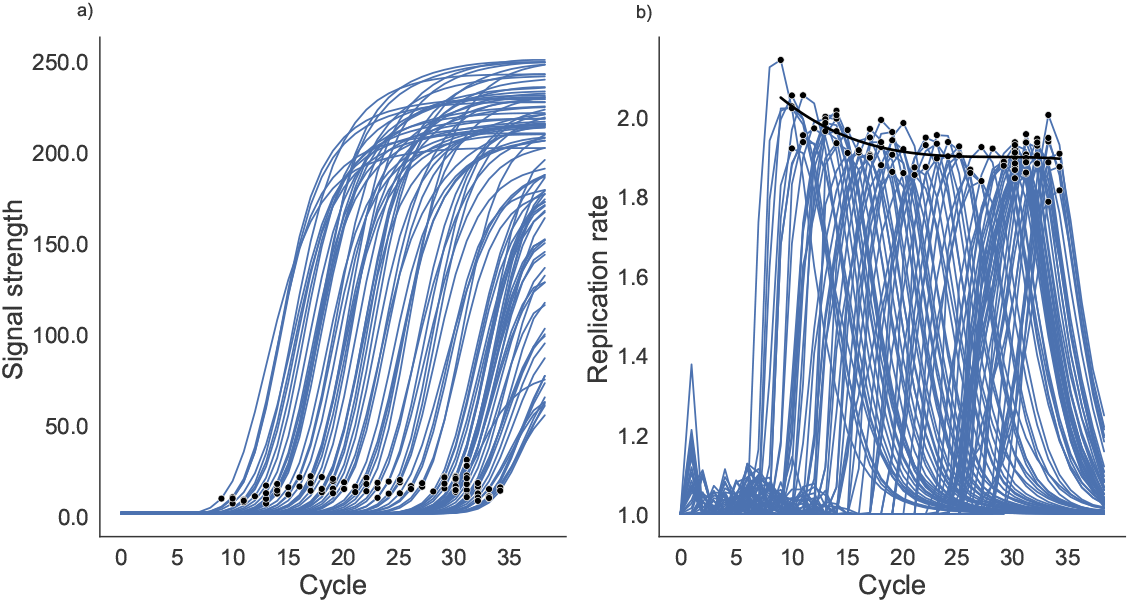
a) PCR traces and the points at the maximum replication rate (black dots). b) the replication rate for each trace with the points where maximum replication rate is reached. Notice that the max replication rate falls as cycle increases.

Maximum replication rate is then calculated as the largest ratio of the signal at a given cycle to the signal at the previous cycle. A polynomial regression is fit to the relationship between maximum replication rate and cycle at maximum replication rate, yielding *β*. The degree of the polynomial is chosen via a cross-validation grid search over degrees 1, 2, 3 (linear, quadratic, and cubic), providing a more robust update to the description Arnaout 2021. With *β* estimated and *v*_*L*_ and 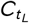 provided by the user, the integrals in equation (1) are calculated numerically by *ct2vl*. Equation (1) is now calibrated and is ready to convert *C*_*t*_ values to viral loads. The user is spared interaction with the mathematics (see Usage, below).

### Calibration and validation datasets

The FDA-approved Abbott Alinity m Real-Time PCR assay SARS-CoV-2 RT-qPCR testing platform was used for this analysis. Results for the Abbott m2000 have been previously described (12). To validate *ct2vl*’s accuracy, we compared the viral load predictions from *ct2vl* to a validation dataset composed of 40 *C*_*t*_ values and their corresponding viral loads from two independent calibration series of viral loads on the same Abbott Alinity m SARS-CoV-2 RT-qPCR machines. First, we calibrated *ct2vl* on 96 positive PCR traces from one of the Alinity machines, using equation (1) and the known (experimentally confirmed) *LoD* = 100 copies/mL and mean 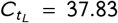 for this machine (see Usage, below). We then used *ct2vl* to convert the 48 *C*_*t*_ values from the validation dataset to viral loads and compared these predicted viral loads to the ground-truth viral loads in the validation dataset to estimate the prediction error. The calibration and validation datasets are provided as Supplementary Information.

For the validation dataset, the genome copy number was based on the reference standard produced by SeraCare (AccuPlex SARS-CoV-2 Reference Material Kit, catalog number 0505-0126). This control material consists of replication-incompetent, enveloped, positive-sense, single-stranded RNA Sindbid virus into which SARS-CoV-2 PCR targets detected by Abbott SARS-CoV-2 RT-qPCR assays have been cloned. This control material was quantified by the manufacturer using digital droplet PCR, and diluted into viral transport medium for analysis.

### Sensitivity analysis

To estimate *ct2vl*’s sensitivity to the 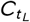 parameter, we replaced the mean value of 37.83 with each of 23 different 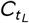 measurements in separate calibration tests (these values were averaged together to get the mean), while holding all other parameters fixed, and measured the prediction error (| predicted viral load - known viral load |) on the validation dataset (for which viral load was known). To estimate *ct2vl*’s sensitivity to *β*, we bootstrap-resampled our calibration data 1000 times (randomly sampling the same number of traces, with replacement), refit the polynomial regression (in Equation (1)) on each bootstrapped sample, then calculated the *ct2vl* prediction error on the validation dataset for each bootstrap sample. To estimate total confidence intervals, i.e., the cumulative effect of variation in *CtL* and *β*, we bootstrap-refit *β* for each of the 23 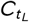 values.

### Code coverage

In computer science, code coverage (or test coverage) is a measure of how much of the code is covered by test suites. Complete coverage means every line of code has been tested for proper function. For *ct2vl*, code test coverage was determined using the *pytest-cov* package.

## RESULTS

### Python package overview

The Python package *ct2vl* takes RT-qPCR-reaction time series as input to parameterize an equation describing the relationship between *C*_*t*_ value and viral load (Equation (1)); after this calibration step, it converts new *C*_*t*_ values to viral load for a given 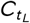 and *v*_*L*_. The package can be used as a command line tool or imported into Python programs. The package has 100% test code coverage and has been tested on macOS, Ubuntu, and Windows.

### Calibration and validation results

For our calibration dataset, the coefficients of the polynomial fit between max replication rate and cycle at max replication rate were 2.27, -2.48e-02, 4.06e-04 (Fig. 3). For the validation dataset, predicted values demonstrated excellent agreement with observed values (Pearson’s *r* = 0.99, *p* < 0.001). The mean absolute error between predicted and observed viral load was 0.21 ± 0.28 log10 units (*mean* ± 2*std*), meaning that predicted viral loads were accurate to within 2.5 ± 1.5 fold, highly accurate considering that viral loads range over 10 orders of magnitude in SARS-CoV-2 infection (12). Consistent with this finding, *R*^2^ was 0.97 for a linear fit between predicted and observed viral loads, with slope 1:1, demonstrating the accuracy as well as precision of *ct2vl* over the full range of the six orders of magnitude of available validation data (Fig. 4).

**FIG 4.**
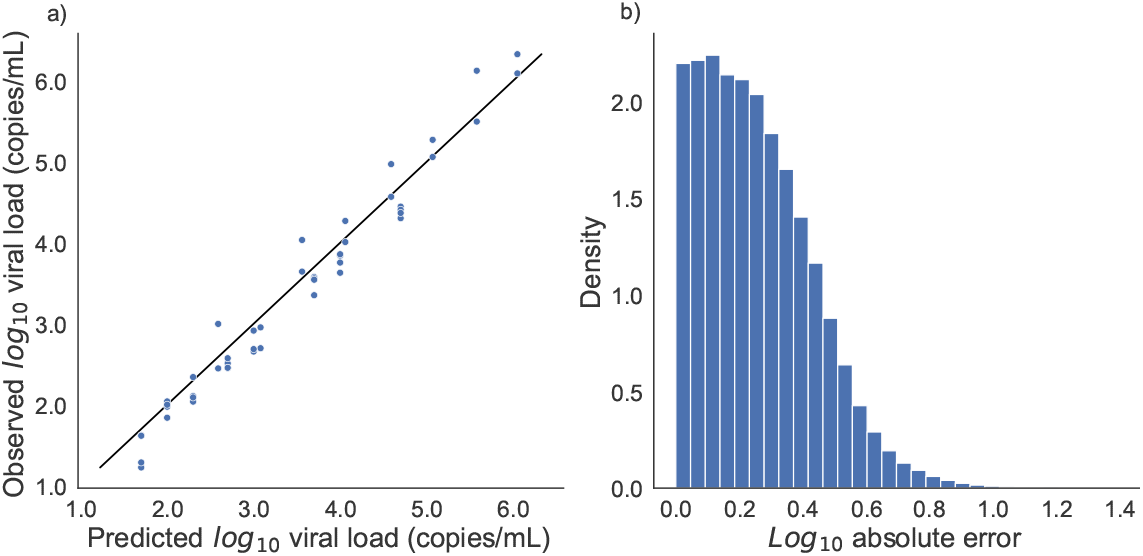
Predicted vs. observed viral load. (a) Viral loads predicted by *ct2vl* compared observed viral loads in a validation dataset of known viral loads (Seracare). (b) Histogram of absolute prediction error. Notice that the majority of the error is below half a log10 unit. Since viral loads can range over 10 orders of magnitudes, this error is within a clinically useful range.

### Sensitivity analysis results

Regarding sensitivity to the 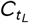 parameter, we found an absolute prediction error of 0.25 ± 0.33 log10 units (*mean* ± 2*std*). Bootstrapping *β* parameters gave a mean absolute error of 0.24 ± 0.30 log10 units. Lastly, to measure overall sensitivity, varying 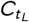 and bootstrapping *β* parameters simultaneously gave a mean absolute error of .25 ± 0.35 log10 units (Fig. 5).

**FIG 5.**
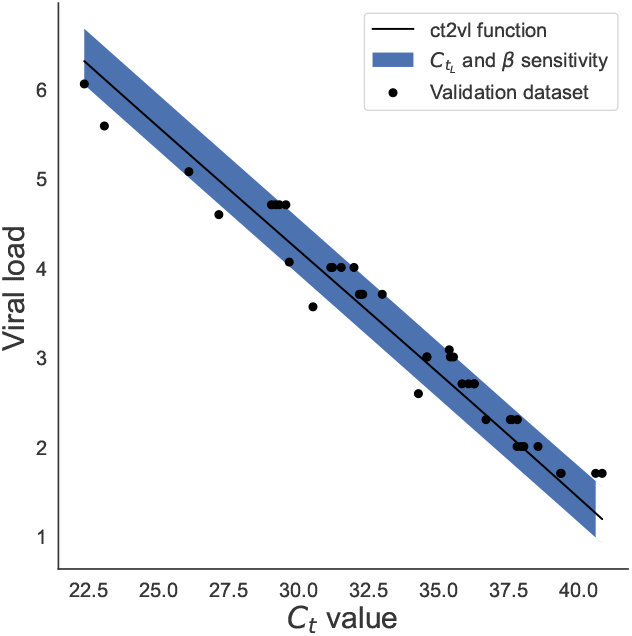
*C*_*t*_ value vs. viral load. The validation dataset (black dots) with the *ct2vl* prediction function (mean ± 2std) when 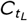 was varied and the calibration parameters (*β*) were bootstrapped.

### Installation

*ct2vl* requires Python 3.7 or higher to be installed. Assuming *pip* is installed, to install *ct2vl*, at the command line, run

~~~
$ pip install ct2vl
~~~

### Command-line usage

To calibrate *ct2vl* run

~~~
$ python3 −m ct2vl calibrate <traces > <LoD> <Ct_at_LoD>
~~~

Here <infile> is a csv file containing the positive traces where each row is a PCR reaction trace and each column is a time step in that trace. See the example file positive_traces.csv in the Supplementary Information, for which the command would be:

~~~
$ python3 −m ct2vl calibrate traces. csv 100.0 37.83
~~~

Once *ct2vl* has been calibrated using the above command, *C*_*t*_ values can be converted to viral loads by typing

~~~
$ python3 −m ct2vl convert <Ct>
~~~

One or multiple *C*_*t*_ values can be passed. For example, to convert a *C*_*t*_ value of 23.1 to a viral load, with a LoD of 100 copies/mL and a corresponding *C*_*t*_ value of 38.73:

~~~
$ python3 −m ct2vl convert 23.1
~~~

The output will be printed to the screen in a text table with integer row numbers, the LoD and Ct-at-LoD 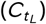 used, the *C*_*t*_ value that was input, the viral load in units of copies/mL, and the viral load in log10 units:

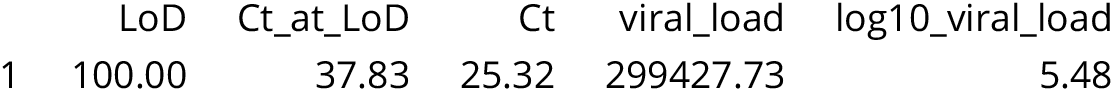

To convert several *C*_*t*_ values to viral loads:

~~~
$ python3 −m ct2vl convert 25.32 30.11 35.95
~~~

Output:

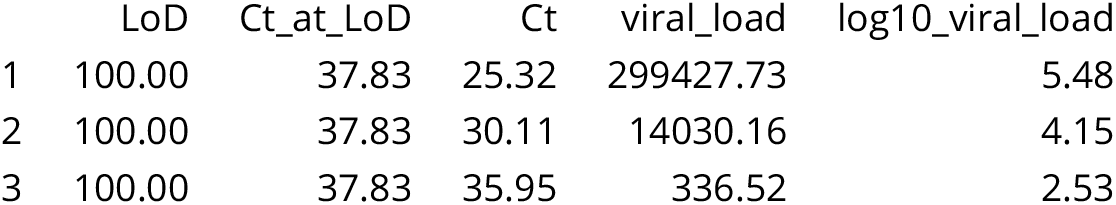

Output can be saved to a file by providing a file path to the optional flag ‘–output’, like so:

~~~
$ python3 −m ct2vl convert 100.0 37.83 23.1 −−output viral_loads.tsv
~~~

Here, the tabular output will have been saved to a tab-delimited text file called viral_loads.tsv.

### Python-package usage

For users who are familiar with the Python programming language and environments, *ct2vl* can also be used programmatically as follows:

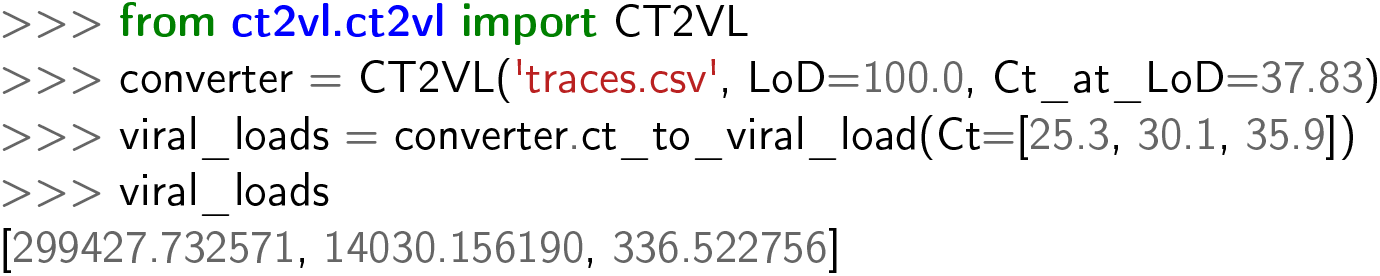

### Availability

*ct2vl* is freely available on the Python Package Index ([PyPI](https://pypi.org/project/ct2vl/)) or via [GitHub](https://github.com/Elliot-D-Hill/ct2vl).

## DISCUSSION

The COVID-19 pandemic necessitated development and approval of RT-qPCR tests at a rate that outstripped the ability to convert results reliably from *C*_*t*_ values to viral loads. As COVID-19 becomes endemic, the probability that patients will receive results on multiple platforms can only rise, as will the need to manage cases where viral load will need to be monitored over time. *ct2vl* facilitates calculation of viral loads for any platform, based on a laboratory’s own validation data. Because the mathematics of RT-qPCR are more complicated for real-world clinical tests, which often contain internal controls and multiple targets, than in stripped-down experimental systems, and because accurate assessment of maximum efficiency is more important for viral load estimation than fitting the entirely of the curve (lag, log, and stationary phase), *ct2vl* concentrates on the most important cause of deviation from pure exponential growth— the fall of replication efficiency with cycle—and fits its empirically/phenomenologically, as opposed to shoe-horning a particular S-shaped curve from the many such curves that exist.

Comparison against a calibration curve of well described SARS-CoV-2 standard (Seracare) demonstrated excellent performance of *ct2vl*, including robustness to sensitivity analysis. Total error was less than half a log10 unit, acceptable performance relative to the 10 log10 units over which viral loads vary in SARS-CoV-2 infection and comparable to the error in HIV viral load testing. *ct2vl* is made available free of charge with a completely open-source codebase and 100% code test coverage, to facilitate customization and incorporation into laboratory workflows. We note that *ct2vl* is applicable to any situation in which calibration is available and polymerase replication rate does not rise with cycle number (i.e., all conventional PCR). It is hoped that it will prove useful beyond SARS-CoV-2, even as the world continues to have to manage successive waves of COVID-19 (16).

## SUPPLEMENTARY INFORMATION

The traces used to calibrate ct2vl for this study and validation data can be found [here](https://gist.github.com/Elliot-D-Hill/1ef5b826d23ffd6f29397958ca23eb7b).

